# Memory in trait macroevolution

**DOI:** 10.1101/465971

**Authors:** Emma E. Goldberg, Jasmine Foo

**Affiliations:** Department of Ecology, Evolution & Behavior; University of Minnesota; Saint Paul, MN; Department of Mathematics; University of Minnesota; Minneapolis, MN November 6, 2018

**Keywords:** comparative methods, trait evolution, phylogenetics, renewal process

## Abstract

The history of a trait within a lineage may influence its future evolutionary trajectory, but macroevolutionary theory of this process is not well developed. For example, consider the simple binary trait of living in cave versus surface habitat. The longer a species has been cave-dwelling, the more may accumulated loss of vision, pigmentation, and defense restrict future adaptation if the species encounters the surface environment. However, the Markov model of discrete trait evolution that is widely adopted in phylogenetics does not allow the rate of cave-to-surface transition to decrease with longer duration as a cave-dweller. Here, we describe three models of evolution that remove this ‘memory-less’ constraint, using a renewal process to generalize beyond the typical Poisson process of discrete trait macroevolution. We then show how the two-state renewal process can be used for inference, and we investigate the potential of phylogenetic comparative data to reveal different influences of trait duration, or ‘memory’ in trait evolution. We hope that such approaches may open new avenues for modeling trait evolution and for broad comparative tests of hypotheses that some traits become entrenched.

## Introduction

One style of studying trait macroevolution is to investigate commonalities in how a trait evolves across diverse lineages. By abstracting away the ecological and evolutionary processes that act on short timescales, a single question can be posed across hundreds of species and millions of years. For example, one big question is whether the evolution of certain traits is irreversible (Bull and Charnov 1985). Existing models of transitions among categorical trait values can test this question on phylogenetic data (Lewis 2001; Nosil and Mooers 2005; Goldberg and Igić 2008), focusing on the emergent pattern of trait evolution asymmetry while sweeping aside details like how it is caused by asymmetry in selective regime shifts or in the capacity to adapt to such shifts. Similarly, phylogenetic comparative methods are available to ask many other questions about trait macroevolution, such as whether traits change more rapidly in some clades than others (O’Meara et al. 2006; Beaulieu et al. 2013), or whether traits tend to change more during speciation than within single lineages (Bokma 2008; Goldberg and Igić 2012; Magnuson-Ford and Otto 2012). Such abstracted models have been very useful, both because they are simple enough to be interpreted broadly and because they can be fit statistically to large phylogenetic datasets. But traits may also evolve in emergent modes that are not captured by existing models. Here, we suggest that a different dynamic of trait evolution may also be widely applicable and mathematically tractable.

Our focal question is, does the length of time a lineage has held a trait value affect the chance of the trait changing in the future? At the macroevolutionary scale, we envision this pattern as the result of two components. In the first component, time spent in one state may lead to increased fit to that state. One possible mechanism is an accumulation of adaptive changes. For example, flowers can become increasingly suited to long-tongued pollinators via gradual elongation of nectar spurs and petal color changes from purple to red to white (Whittall and Hodges 2007). Or focusing on the genetic level, fusions that unite loci determining sex with loci experiencing sexually antagonistic selection can eventually create heteromorphic sex chromosomes in species with separate male and female individuals (Charlesworth 2015). Another possible mechanism is gradual degradation through disuse. For example, vision genes are downregulated in recently-derived cave-dwelling fish populations and accumulate loss-of-function mutations in older cavefish species (Niemiller et al. 2013; McGaugh et al. 2014). In the second component, increased commitment to one state may reduce the chance of changing to another state. In par-ticular, it could take longer to reverse the evolution of more extensive adaptations or losses. This logic seems reasonable and has some theoretical basis (Marshall et al. 1994), but well-supported empirical examples are elusive. For the sex chromosome example above, flowering plant species with heteromorphic sex chromosomes appear less likely to transition back to hermaphroditism than do other dioecious species (Goldberg et al. 2017). For the other examples above, the logic would be that species with longer nectar spurs would be less able to change to short-tongued pollinators when the pollination environment shifted to bees, or cavefishes with more extensive loss of vision and pigmentation would be less able to establish surface populations when washed into aboveground habitats. More broadly, macroevolutionary studies frequently focus on widely-recorded and ecologically-important traits (e.g., diet, habitat, reproductive or life history strategy) that are underlain by an assortment of morphological, physiological, and behavioral attributes with complex genetic bases. If these attributes accumulate gradually and inhibit subsequent changes in the focal trait, it may be common for the history of a trait within a lineage to affect its propensity for evolutionary change in the future.

Although it seems intuitively reasonable that a lineage’s duration in one state could affect the chance of change to another state, this dynamic is absent from the model that dominates phylogenetic studies of discrete trait evolution. In the existing model, evolutionary changes between states occur as jumps with specified probabilities (Pagel 1994; Lewis 2001). Variations on the theme are numerous. State space can be structured to accommodate everything from codons to geographic ranges to correlations between multiple traits, rates of state change can depend on time or clade, and trait evolution can interact with the speciation-extinction process (Felsenstein 1981; Goldman and Yang 1994; Pagel 1994; Ree et al. 2005; Maddison et al. 2007). One core assumption remains throughout all these variants, however: the length of time that a lineage has possessed its state does not affect the probability that it will change state. That is, these are all ‘memory-less’ Markov models. Recent non-Markovian models for lineage diversification allow the age of a lineage to influence its probabilities of speciation or extinction (Stadler 2013; Hagen et al. 2015; Alexander et al. 2016). For trait evolution, however, the only previous non-Markovian model is the threshold model (Felsenstein 2005), which we discuss in detail below.

Here, we present models that incorporate the dynamic of ‘memory’ in trait macroevolution. We retain the abstract simplicity of representing evolution as jumps between discrete states, but we add the possibility that these jumps are affected by how long a lineage has held its state. First we derive mathematical forms for the memory dynamic from simple assumptions about its underlying cause. Then we investigate whether phylogenetic comparative data can reveal the signature of memory in trait macroevolution. We close by discussing how future work could further open this macroevolutionary idea to empirical study.

## Models

### Renewal process

For modeling the evolution of discrete-valued traits on a phylogeny, a continuous-time Markov chain is by far the most common approach (Felsenstein 1981; Pagel 1994; Lewis 2001). In this model, the chance of a change in state depends only on the rate parameters and the current value of the state. For example, if the trait can take either state *A* or *B*, the model is described by two parameters: *q_AB_* is the instantaneous rate at which a lineage in state *A* flips to state *B*, and *q_BA_* is the instantaneous rate for the reverse trait flip. (Throughout, we will consider only binary traits, so a ‘flip’ is a change to the other state.) The trait flips from state *A* to *B* follow a Poisson process in this model, and the waiting time until the next flip has an exponential probability distribution with mean 1/*q_AB_* (and similarly for flips from *B* to *A*).

Our goal is to build a model in which the instantaneous rate of a trait flipping depends on how long the lineage has held that state. This requires removing the ‘memory-less’ property of the Markov and Poisson processes, rendering the waiting times no longer exponentially distributed. The renewal process is the generalization of the Poisson process to any distribution of waiting times, provided they are still independent and identically distributed (Ross 2010, Ch. 7). Each trait flip constitutes a ‘renewal,’ and the time until the next flip depends on the time since the last renewal. Our derivations will consider only the symmetric case in which transitions from *A* to *B* have the same distribution as from *B* to *A*. Future work could relax this assumption by using an alternating renewal process.

The ‘hazard function’ describes the instantaneous rate of an event occurring. In our context, this is the chance of a flip occurring at time *t* given that the previous flip was at time 0 (fig. 1). In terms of the probability density function (PDF) of the waiting times, *f* (*t*), and its cumulative distribution function (CDF), *F*(*t*), the hazard function is *h*(*t*) = *f* (*t*)/[1 − *F*(*t*)]. For the usual Poisson process of trait flips, the hazard function is flat, e.g., *h*(*t*) = *q_AB_*. Under the idea that extended commitment to one state inhibits evolutionary transitions to another state, we would like a trait evolution model with a declining hazard function, so *h*(*t*) decreases with *t*. There could perhaps be other situations in which an increasing hazard function is appropriate, and our derivations also allow for this. For example, a parasite may be more likely to switch hosts after enough time has passed that its current host has adapted to reduce its efficacy.

**Figure 1:**
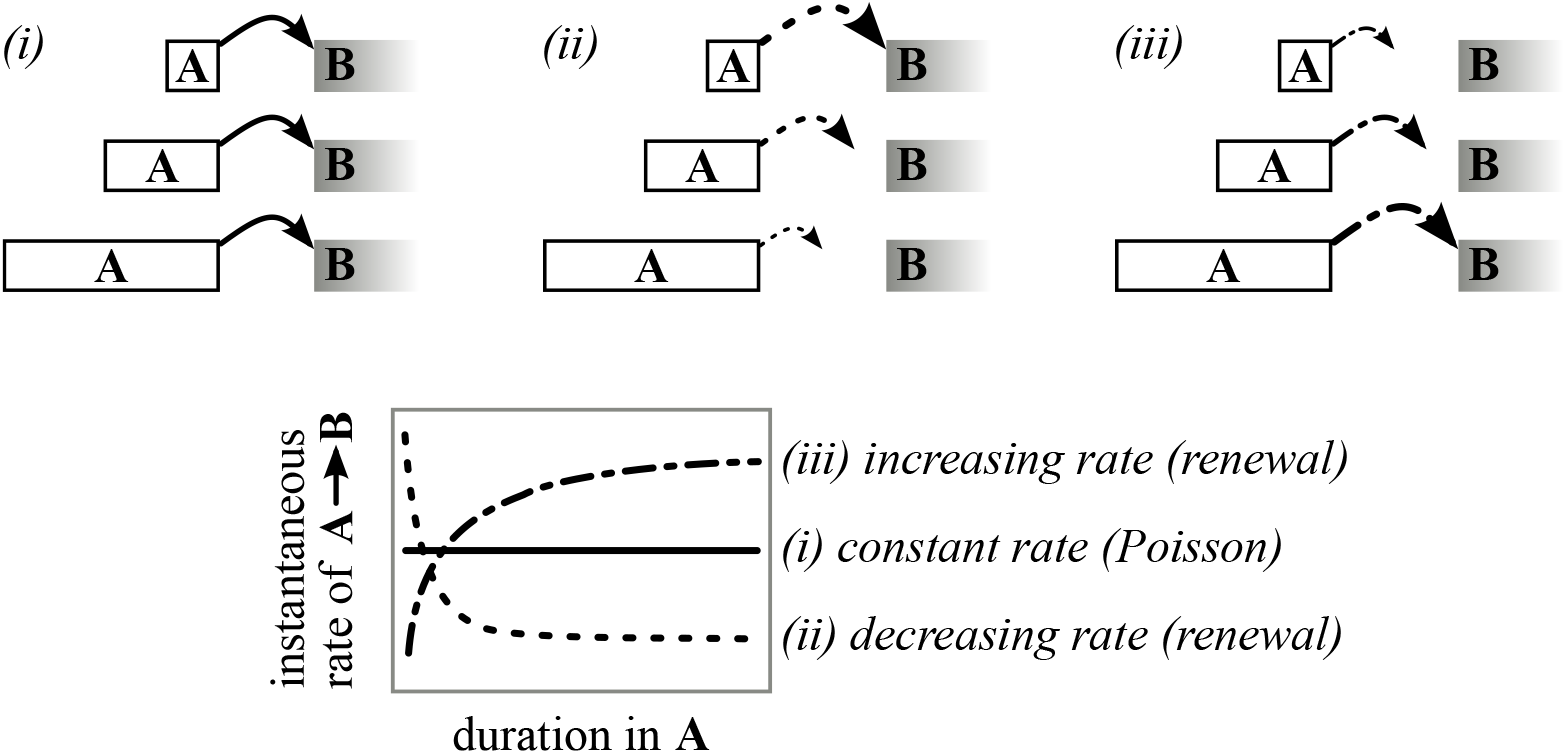
Transitions from state *A* to state *B* may be (i) independent of how long a lineage has held state *A*, (ii) less likely as *A* has been held for longer, or perhaps (iii) more likely as *A* has been held for longer. Possible corresponding hazard functions are shown in the lower panel. These are hazard functions of the Gamma distribution, which is specified by ‘shape’ and ‘rate’ parameters. The hazard is (i) flat when shape = 1, (ii) decreasing when 0 < shape < 1, or (iii) increasing when shape > 1. The rate parameter is the value after a very long duration in *A*.

The renewal process in general can operate with any hazard function. What is an appropriate renewal function for trait evolution? We next describe three models that abstract the process of trait evolution with different forms of ‘memory.’ We derive the hazard function for each and then compare across models.

### Threshold models

There is currently one phylogenetic model of discrete trait evolution that inherently causes the duration in one state to affect the chance of flipping to the other state: the Threshold model (Felsenstein 2005). This model tracks the evolution of an unobserved continuous-valued quantity called the ‘liability.’ The observed discrete-valued trait takes state *A* when the liability is below a certain threshold value and state *B* when it is above the threshold (fig. 2A). This model represents the situation in which a trait can only take discrete observable states, such as presence or absence, but a large number of genetic and environmental factors together determine the state (Wright 1934).

**Figure 2:**
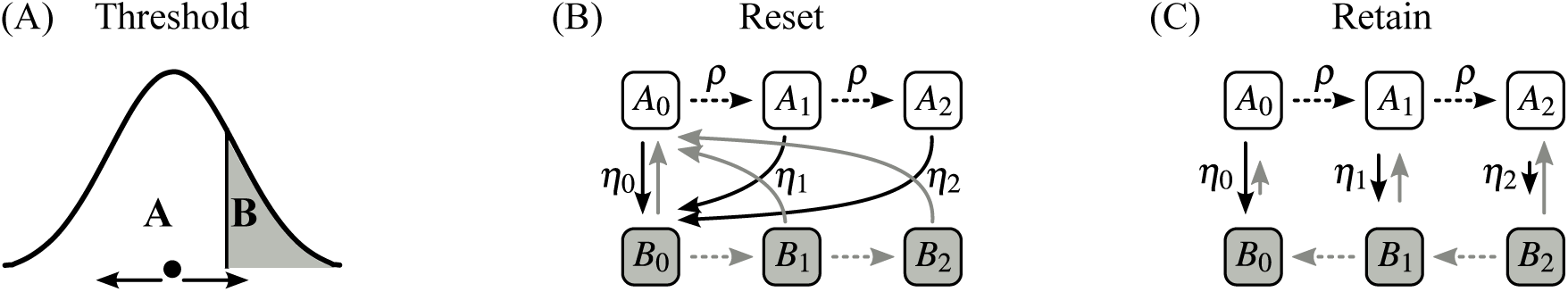
Three models for the evolution of a trait that can take observable states *A* or *B*. (A) In the Threshold model, a liability value evolves on a continuous scale, and the corresponding discrete state is determined by whether the liability is less than or greater than a threshold value. (B) In the Reset model, changes accrue while a lineage holds a state, and flips to the other state always reset the value to the corresponding initial substate (*A*_0_ or *B*_0_). (C) In the Retain model, changes also accrue but in opposite directions for each state, and the substate value is retained upon transition to the other observed state. In (B) and (C), dashed arrows show transitions between unobserved substates (with rates *ρ*) and solid arrows show flips to the other observed state (with rates *η_i_*).

It is intuitive that memory is built into the evolution of such a trait. The longer the state has remained *A*, the farther is the liability expected to have wandered from the threshold, making a transition to *B* less likely. The Threshold model has been used to compute correlations between traits (Felsenstein 2005) and to infer ancestral states (Revell 2014). Here we relate the Threshold model to a renewal process of trait evolution to better understand its memory properties.

The original threshold trait model describes normally-distributed liability values (Wright 1934), and a Brownian motion process was later used for the evolution of the liability (Felsenstein 2005). The Brownian motion formulation is, however, not suited to our goal of modeling the time to the next trait flip. If *t*_1_ is the time at which the trait value crosses the threshold into a particular state and *t*_2_ is the next time that the trait returns to the threshold and flips back to the previous state, then for any *ϵ* > 0 we have that *P*(*t*_2_ < *t*_1_ + *ϵ*) = 1. That is, the probability of returning to the threshold is one even over a vanishingly small amount of time. Thus, although the Brownian motion formulation used by Felsenstein (2005) works well for other applications of the Threshold model, we need an alternative formulation to compute a meaningful distribution of times until the next trait flip.

#### Random walk model

We describe a different model for the liability, which retains the spirit of the Threshold model but avoids the artificial pathological path properties of Brownian motion. Consider a one-dimensional random walk in which steps of size one to the left or the right are equally likely, and the waiting time between steps is exponentially distributed with rate *θ*. For convenience, we place the threshold at 0.5: the trait thus flips from *A* to *B* when the liability steps from 0 to 1, vice versa for the other direction, and the liability spends no time directly on the threshold.

We are interested in the probability distribution of *τ*, the amount of time it takes to flip to *B* if *A* has just been acquired. (It is the same for flips in the reverse direction because our random walk is symmetric, but we pick one case for clarity.) Let *f_τ_* and *F_τ_* be the PDF and CDF, respectively, of *τ*. Let *N* be the number of steps taken by the random walk before hitting 1 for the first time, starting from 0; this is the number of steps between threshold crossings. Then for positive integers *i*, the probability mass function of *N* is given by (Lalley 2016)

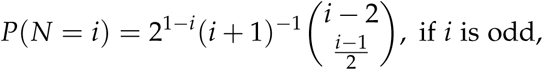

and *P* (*N* = *i*) = 0 for all even values of *i* due to the parity of the random walk.

The times between steps of our random walk are exponentially distributed with rate *θ*, so the time *τ* can be interpreted as a sum of *N* independent exponential random variables each with rate *θ*, where *N* is itself a random variable. The sum of independent identical exponential random variables has a Gamma distribution (Ross 2010, Ch. 5). Therefore, conditioned on *N* taking some particular value *i*, the distribution of time to the next flip is *τ* = *Y_i_* where *Y_i_* is a Gamma random variable with shape parameter *i* and rate parameter *θ*. Allowing for all possible values of *N*, we can then write the PDF or CDF of *τ* as a mixture of PDFs or CDFs of the *Y_i_*, for *i* = 1, 2, …. The hazard function of *τ* thus becomes

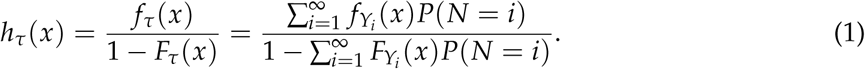

The hazard function for the symmetric random walk Threshold model (eq. [1]) is illustrated in figure 3A. The rate of flips to state *B* always decreases with time spent in *A*. The steepness of that decrease is determined by the distribution of times between steps. With larger values of *θ*, the time between steps is smaller, so the liability quickly wanders farther from the threshold and a flip to the other state rapidly becomes less likely. When the time spent in *A* is longer, the random walk is more likely to have already wandered far from its starting point, so waiting additional time does not significantly affect the rate of flipping to *B*. In this regime, the dependence on *θ* also decreases due to the following compensatory mechanism: for fixed time, larger values of *θ* result in the walk being farther from the threshold, requiring more steps to return taken at a faster rate, while smaller values of *θ* are associated with the walk being closer to the threshold, requiring fewer steps to return but taken at a slower rate.

**Figure 3:**
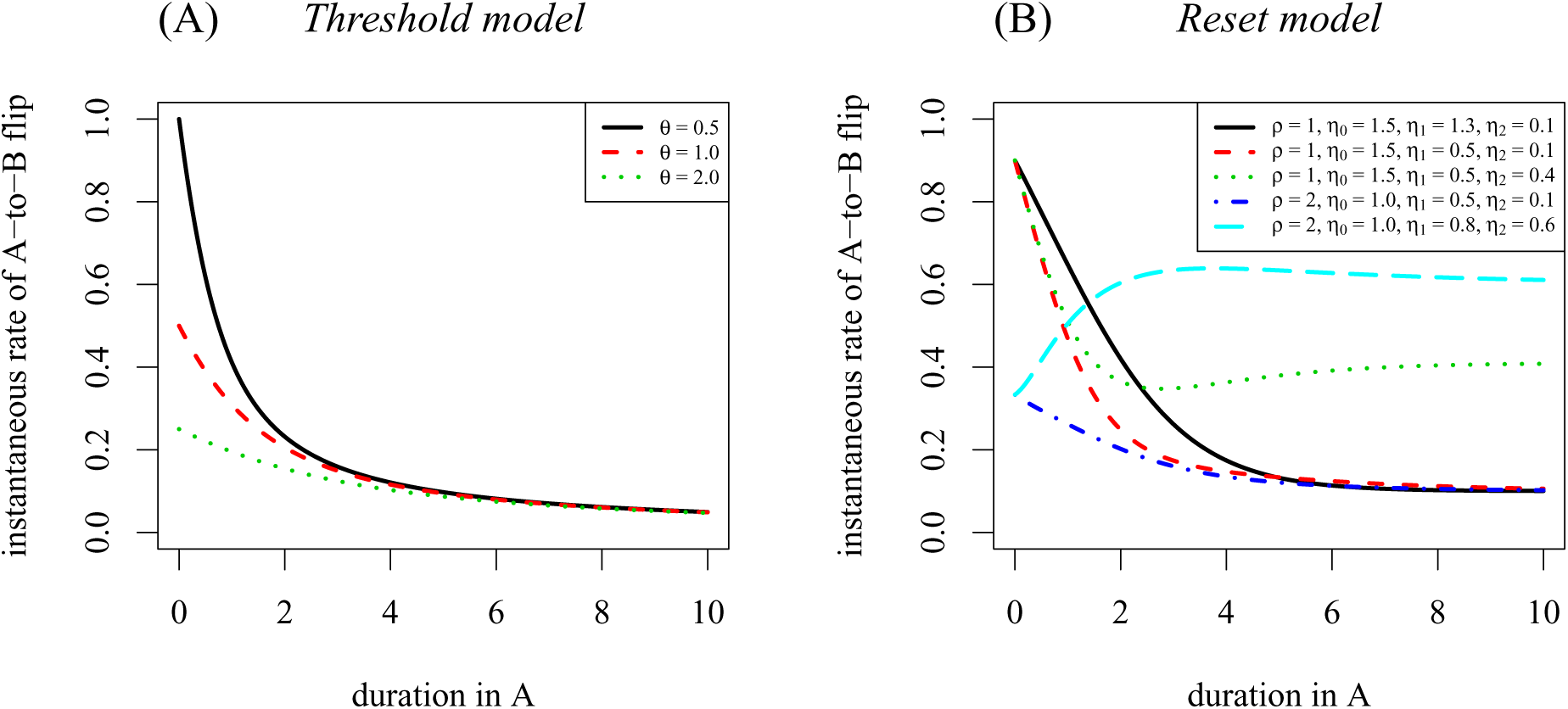
Hazard functions for the Threshold and Reset models. (A) In the symmetric random walk threshold model (fig. 2A), the rate of flips to state *B* always decreases with time spent in *A* (eq. [1]). Larger values of *θ* correspond to less time between steps, so the liability more quickly wanders away from the threshold. (B) In the model where the subtraits are reset upon a flip to the other state (fig. 2B), a variety of hazard function shapes are possible even when *η*_0_ > *η*_1_ > *η*_2_ (eq. [2]). In many situations, the rate of flips to state *B* decreases with time spent in *A*. But when progression within a state (at rate *ρ*) outpaces flips to the other state (rates *η_i_*), the dynamics can be drawn into a longer path from *A* to *B*.

### Multi-state models

Another way to conceptualize a process that produces memory in trait evolution is an accumulation of changes in other traits (‘subtraits’) that support the focal trait. For example, if the focal trait is diet type, a species may become increasingly more adapted to eating insects as it acquires the behavioral, morphological, and physiological attributes that allow it to find, catch, and digest that type of prey. Alternatively, the subtraits could represent accumulated losses of function in genes that are no longer under selection, such as functional eyes or pigmentation once a species becomes cave-dwelling. Even if it would be possible to observe these subtraits, perhaps not all have been identified or included in a dataset focused on the main trait of interest. We will therefore assume that only the focal trait, with values *A* or *B*, is observed, and not the values of the subtraits (called *A_i_* and *B_i_* for *i* = 0, 1, …).

Structured multi-state Markov models have previously been used to describe the macroevolution of subtraits within focal traits. For example, Zenil-Ferguson et al. (2017) considered transitions between two states, herbaceous and woody, while simultaneously modeling changes in chromosome number within each state. All the modeled states are observable in this case, because they are combinations of growth form and chromosome number. In contrast, Beaulieu and O’Meara (2016) add a hidden state to a model of binary trait evolution, so that each observed state is represented as two hidden substates between which transitions are possible. Applying this model to plant breeding systems, Freyman and Höhna (in review) found the hidden state to represent a memory process: lineages evolved from *A* to one hidden state of *B* and then to the other hidden state of *B*. (The hidden states were indistinguishable phenotypically, but they had different effects on lineage diversification.) Tarasov (in review) describes other arrangements of multi-state Markov models for the evolution of traits with hidden or hierarchical aspects.

We next describe two multi-state models explicitly structured to represent memory in trait evolution (fig. 2BC). In each, we assume that as time passes, a lineage evolves through a sequence of substates that underly the focal trait. In the examples mentioned above, this could represent increasing adaptation to an insectivore diet or increasing loss of function within a cave environment. Both of our multi-state models exhibit memory when the rate of flipping to the other focal state depends on the current substate. The two models differ in the effect that a flip in the focal trait has on the value of the subtrait. In the Reset model (fig. 2B), the subtrait value that accumulated in the previous focal state is reset because it is irrelevant when that focal trait changes. For example, progression through insectivore subtraits might involve gradually gaining the ability to distinguish palatable from noxious insect prey, but this subtrait may have no cost or benefit when the predominant food changes to seeds. In the Retain model (fig. 2C), the subtrait value that accumulated in the previous focal state is retained and thus has an immediate effect when the focal trait changes. For example, progression through cave subtraits might involve gradually losing functional eyes, and that reduced vision would still be present in a lineage that just transitioned to surface habitat. We explain each model further below, but in essence the distinction is whether increased entrenchment in one focal state is undone immediately or gradually upon transition to the other state. Real traits might exhibit some mix of these two dynamics, but it is informative to consider their separate effects. For each model, we derive their hazard functions in order to compare their memory properties.

#### Reset model

We first consider the case where a flip to the other observed state causes the unobserved subtrait to ‘reset’ its values. Consider a small example with three subtraits (fig. 2B; though our derivation can easily be generalized to more subtraits). Suppose that progressive commitment to *A* is represented as transitions from *A*_0_ to *A*_1_ to *A*_2_, each taking place after an exponentially-distributed waiting time with rate *ρ*. From any of these substates *A_i_*, the species may transition to the first substate of the other observed state, *B*_0_. We assume that these flips also have exponential waiting times, but with rates *η_i_* that depend on the initial substate, *i* = 0, 1, 2. Thus, when *η*_0_ > *η*_1_ > *η*_2_, lineages that have progressed to later substates (*A_i_* for larger *i*) are less likely to flip to state *B*. Our goal is to determine the distribution of *τ*, the time it takes to flip to *B* after entering *A*. In this Reset model, *τ* describes the time to enter *B*_0_ after having just arrived in *A*_0_. (Our symmetry assumptions ensure the answer is the same for flips from *B* to *A*.)

To derive the distribution of *τ*, we consider all the possible paths a lineage could take from *A*_0_ to *B*_0_. For three substates, these are: *A*_0_ → *B*_0_, *A*_0_ → *A*_1_ → *B*_0_, and *A*_0_ → *A*_1_ → *A*_2_ → *B*_0_. Define the random variable *Y* as the substate of *A* just before the flip to *B*. For the three paths above, *Y* = 0, 1, or 2, respectively. In addition, define independent random variables for the transition time to the next substate, *Z_i_* ~ exp(*ρ*) (for *i* = 1, 2), and for the next flip to the other state, *Q_i_* ~ exp(*η_i_*) (for *i* = 0, 1, 2). Then we can rewrite *τ* in terms of these random variables, conditioned on *Y*:

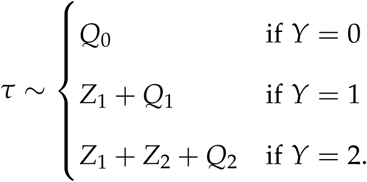

We next define random variables representing renewal times for each of the possible paths: *D*_0_ ≡ *Q*_0_, *D*_1_ ≡ *Z*_1_ + *Q*_1_, *D*_2_ ≡ *Z*_1_ + *Z*_2_ + *Q*_2_. Then we obtain the PDF and CDF of each *D_i_*:

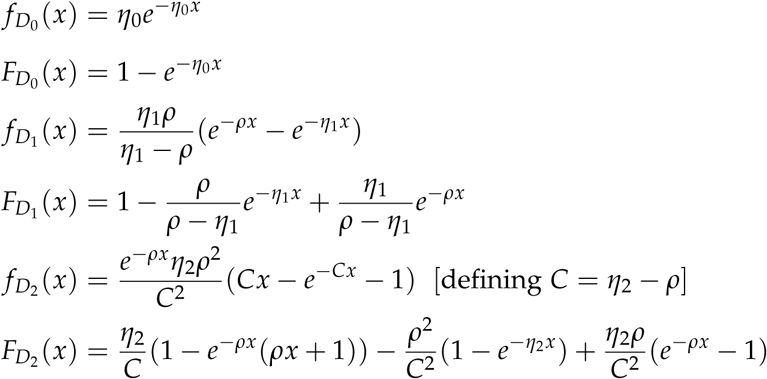

provided *η*_1_ ≠ *ρ*; otherwise *D*_1_ is distributed as a Gamma random variable with shape 2 and rate *ρ*.

In addition to the above expressions for the renewal time along each possible path, we need to know how likely it is to take each path. The conditioning probabilities are the probabilities of each path from *A*_0_ to *B*_0_, i.e., the probabilities that *Y* = *i*:

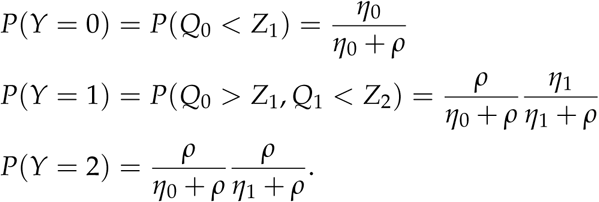

The PDF and CDF of *τ* are then obtained as the distributions for each possible path weighted by the probability of taking that path,

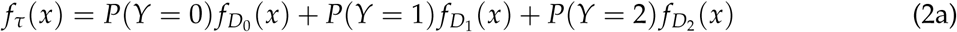

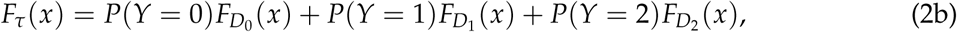

from which we obtain the hazard function,

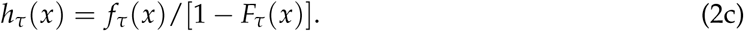

Examples of the hazard function for the Retain model (eq. [2]) are illustrated in figure 3B. A variety of hazard function shapes are possible even when it is increasingly hard to leave subsequent substates (*η*_0_ > *η*_1_ > *η*_2_). When no time has passed in *A*, the rate of flipping to *B* is always *η*_0_/(*η*_0_ + *ρ*), which is the probability of transitioning to *B* rather than to *A*_1_. When a long time has passed in *A*, the rate of flipping to *B* is always *η*_2_ because there is no other option for a transition out of *A*_2_ (in this example with only three subtraits). For intermediate durations in *A*, the shape is determined by the weighted contributions of each possible path to *B*. This allows for hazard functions that are not monotonically decreasing. This may be surprising at first, but recall that the duration in *A* is influenced not only by the time to transition from *A*_0_ to *A*_1_ and so on, but also by the time to flip to *B*, and the combined effect may not be entirely intuitively obvious. For example, an initial increase in the hazard function results if the rate of progressing within the current observed state (from *A*_0_ to *A*_1_, with rate *ρ*) is higher than the rate of flipping to the other observed state (from *A*_0_ to *B*_0_, with rate *η*_0_), because the dynamics can be initially drawn into a longer overall path from *A* to *B* by first taking a step to *A*_1_.

#### Retain model

We next consider the case where a species ‘retains’ the value of its subtrait when flipping to the other observed state. In contrast to the Threshold and Reset conceptualizations of memory in trait evolution, this Retain model cannot be described by a two-state renewal process. Instead, a different renewal process is needed for each substate. To see this, consider again the example with three subtrait values (fig. 2C). As before, transitions to successive substates (*A_i_* → *A*_*i*+1_) take place after an exponential waiting time with rate *ρ*. In contrast to the Reset model, in the Retain model *A_i_* transitions to *B_i_* instead of to *B*_0_ for *i* = 0, 1, 2, so the lineage retains the *A*-adapted subtraits even after the transition to *B*. Again, these flips from *A_i_* to *B_i_* take place after an exponential amount of time with rate *η_i_*, and *η*_0_ > *η*_1_ > *η*_2_ if flips to *B* become increasingly difficult with greater commitment to *A*. (Because subtrait evolution while in *B* undoes changes accrued while in *A*, we might wish to label and order the rates *η_i_* differently for flips from *B* to *A*, as indicated by the gray arrows in fig. 2C.)

In the Retain model, let *τ_i_* be the the time it takes to flip to *B*, starting from state *A_i_*. For the starting state of *A*_0_, *τ*_0_ has the same distribution as the renewal time in the Reset model (eq. [2]). However, *τ*_1_ has a different distribution. Recall the random variable *Y* which tracks the substate at the time of the trait flip. When the initial state is *A*_1_, *Y* can only take values 1 or 2, so *τ*_1_ can be written as

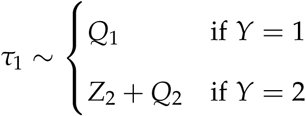

with conditioning probabilities

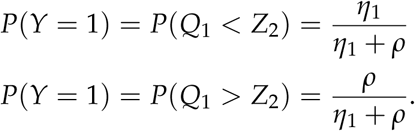

Then we have the PDF and CDF of *τ*_1_:

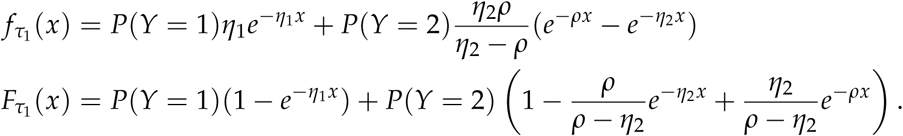

Lastly, *τ*_2_ is simply an exponential random variable with rate *η*_2_, with the corresponding constant hazard function.

Because the renewal time for flips from *A* to *B* depends on the substate held upon arrival into *A*, the renewal process must be modified to explicitly account for all the substates. A two-state renewal process will not suffice. There is thus no single hazard function that describes flips between *A* and *B* in the Retain model. For example, in figure 4 we see that the hazard functions for arrival in *A*_0_ match those of the Reset model with the same parameters (comparing fig. 4A with fig. 3B), but that for those same parameters, the hazard functions for arrival in *A*_1_ and *A*_2_ are different. Similar to the Reset model, as the duration in *A* increases, it becomes more likely that the flip to *B* will occur from the last substate, *A*_2_, so the hazard rates all approach *η*_2_.

**Figure 4:**
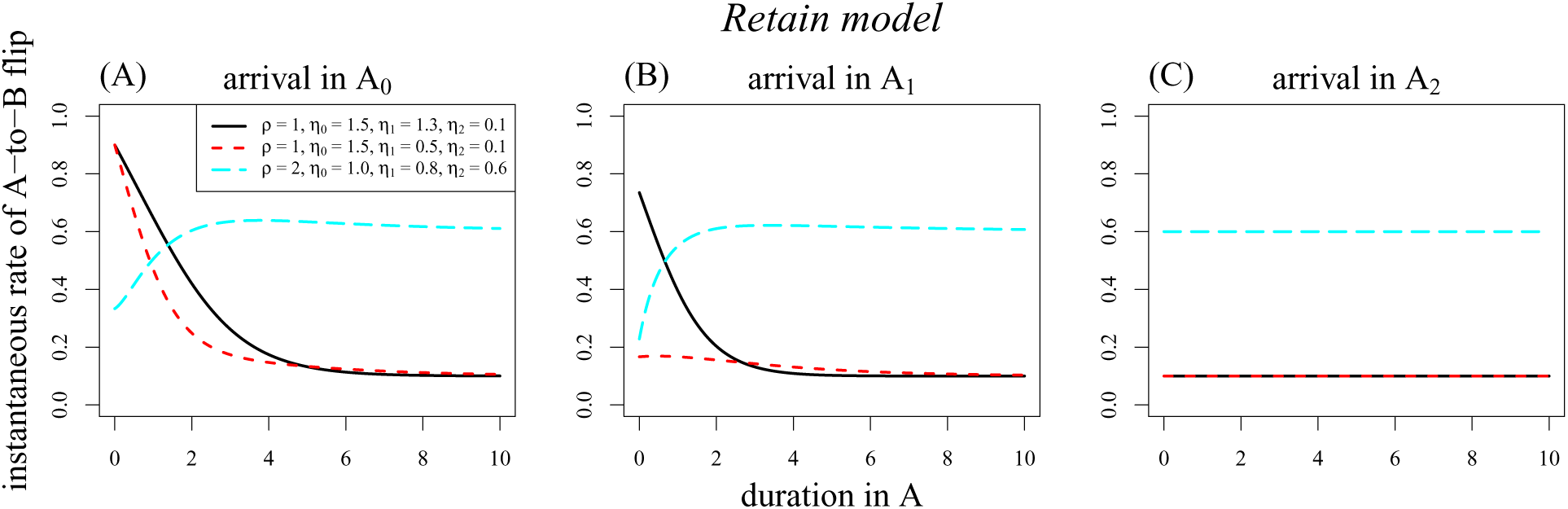
Hazard functions for the Retain model. For the scenario in which adaptation subtraits are retained upon a flip in the focal trait (fig. 2C), the rate of flips to state *B* depends on whether the initial substate was *A*_0_, *A*_1_, or *A*_2_ (panels A, B, and C, respectively). This precludes the use of a two-state renewal process framework.

### Choice of renewal function

All three models considered above contain the idea that changes in many unobserved components accumulate to inhibit changes in the focal binary trait. Each model represents this process differently, however, and we found that the effect is not always the same. The most consistent outcome is a hazard function that declines steeply at first and then more gradually, so that the effect of memory on trait evolution is strongest shortly after a trait change. This is true always for the Threshold model, but only sometimes for the Reset and Retain models. In these latter models, even if the rate of flipping from *A* to *B* declines as subtrait changes accumulate, the hazard function itself need not be strictly decreasing. The Reset model could be fit to phylogenetic data with existing multi-state Markov methods. If this is done, however, our results show that finding *η_i_* > *η*_*i*+1_ (for *i* = 0, 1, …) would be insufficient to conclude that the rate of flips to the other state simply declines with duration in the state.

The above models provide a sense of what a hazard function should look like to be consistent with some abstract mechanisms for how memory may enter trait evolution. Rather than model such mechanisms, however, one could instead work simply with a two-state renewal process and directly specify the mathematical form of the renewal function. This approach would not capture the Retain model, as explained above. However, choosing, say, a Gamma distribution for the renewal function would roughly capture the shape of the hazard seen under the Threshold model and many cases of the Reset model. It also includes as a special case the Poisson model with exponentially-distributed waiting times. Examples are shown in figure 1. We take this approach of directly specifying the renewal function in the next section, when we turn to fitting the renewal process to data.

## Inference

We now consider the question of whether memory in trait evolution can be inferred from phylogenetic comparative data. First, we derive the likelihood of tip character states given the tree and a renewal model of trait evolution. Then, we present a small set of simulation results to test the efficacy of this approach. That is, we investigate whether a Poisson process can be distinguished from a more general renewal process for trait evolution based on commonly-available phylogenetic data.

### Likelihood

To calculate the likelihood of observed tip states on a phylogeny, we employ the pruning algorithm (Felsenstein 1981). Working from the tips of the tree toward the root, this algorithm combines the probabilities of state changes along each branch while summing over possible states at each node. For any model using this algorithm, the key quantity is the transition probability function. Given that a lineage is in state *s*_0_ at time *t*, the transition probability 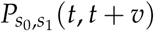 is the probability that the lineage is in state *s*_1_ at time *t* + *v*. We next derive this transition probability for the renewal model.

Our derivation assumes that there are two possible states, and that transitions between them are governed by the same renewal process in each direction. We further assume that we specify directly the renewal function, with PDF *f* and CDF *F*.

To begin, suppose a renewal occurs right at time *t*, creating state *s*_0_ (fig. 5A). The probability of ending up in state *s*_1_ at *v* units of time later is

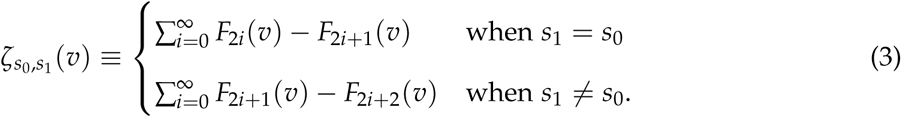

**Figure 5:**
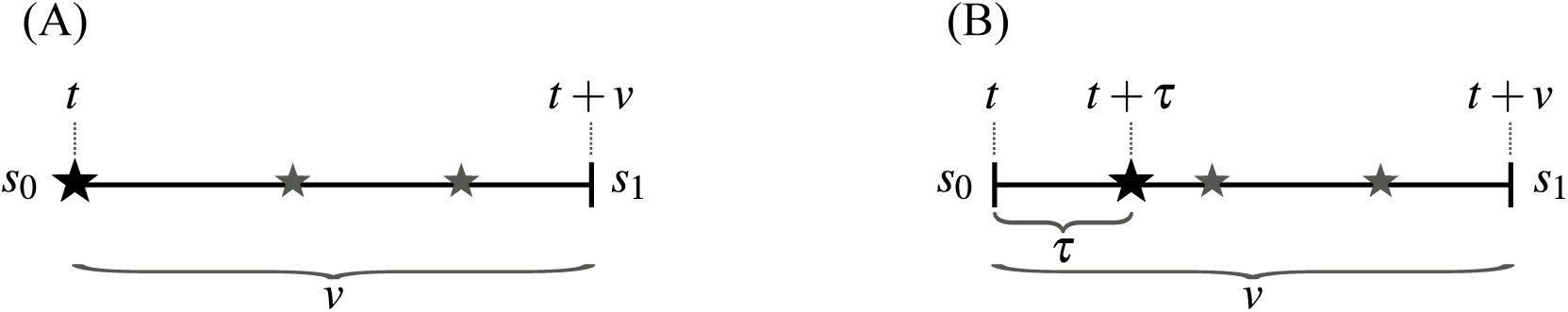
Renewals on a single lineage, used to compute transition probabilities. The initial state is *s*_0_ and the final state is *s*_1_. Renewals are labeled with stars, large and black for the focal event, and small and gray for subsequent events that may or may not occur.

The first case describes an even number of flips during that time, and the second case describes an odd number of flips. The following property of the renewal process is used in equation (3): If a renewal occurs at time 0, let *N*(*t*) be the number of renewals until time *t*. Then *P*(*N* (*t*) = *n*) = *F_n_*(*t*) − *F*_*n*+1_(*t*), where *F_n_*(*t*) is the CDF for the sum of *n* independent copies of the renewal process (Ross 2010, eq. 7.3). That is, *F_n_*(*t*) is the probability that *n* or more renewals have occurred by time *t*, and it is the *n*-fold convolution of *F* with itself. (Note that this convolution is trivial for the Gamma distribution, which is another reason we suggested above that it could be used as the renewal function.)

However, it is in general not the case that a renewal occurs right at time *t*. Let *τ* be the amount of time elapsed from *t* to the next renewal; this is the residual time (fig. 5B). The PDF of *τ* is given

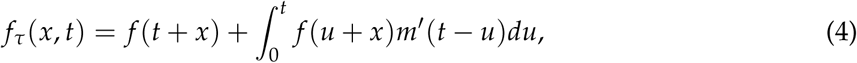

where *m*(*t*) = 𝔼[*N*(*t*)] is the expected value, and *m’* (*t*) = *dm/dt* is the probability that there was a renewal between times *t* and *t* + *dt*. In equation (4), the first term applies when no renewal has happened at all (since time 0), and the second term applies when there was a previous renewal (at time *t* – *u*). This second term integrates over all times that previous renewal could have happened, weighting each by the probability of a renewal then.

If we assume that the trait evolution process is in the limiting regime, we can simplify equation (4):

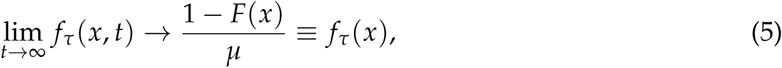

where *μ* is the mean of the distribution *F*. Under this limit, the first term in equation (4) goes to zero because at least one renewal would have happened by *t*. Also, the density of renewal events, *m’* (*t*), goes to its mean value of 1/*μ*, the reciprocal of the mean time between renewals. Thus, we have dropped the dependence on the absolute time *t*, so that *f_τ_* can be interpreted as the amount of time we wait until the next renewal, regardless of the current time. In the following we will retain the assumption that we are concerned only with the limiting regime *t* → ∞, which means assuming that the trait evolution process has run for a long time before the root of the tree.

We now construct the transition probabilities. One possibility is that the first renewal after time *t* occurs before or at time *t* + *v* (fig. 5B). In this case, we must also consider subsequent renewals that may or may not occur by *t* + *v*. Then, the probability of observing state *s*_1_ at time *t* + *v*, conditioned on knowing *s*_0_ at time *t*, is given by:

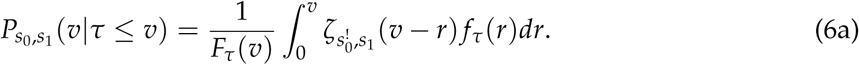

The notation 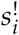 means the state that is not *s_i_*, and *F_τ_* is the CDF of *τ*. We have dropped the *t* dependence from the above equation based on the limiting approximation of the PDF of *τ* (eq. [5]).

The other possibility is that the first renewal after time *t* happens after time *t* + *v*. Then,

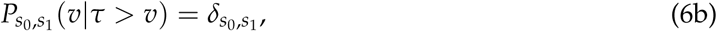

where the Kronecker *δ* function is 1 if the states are equal and 0 otherwise.

Putting these two possibilities (eq. [6]) together, the probability of observing state *s*_1_ at *v* units of time after observing *s*_0_ is given by:

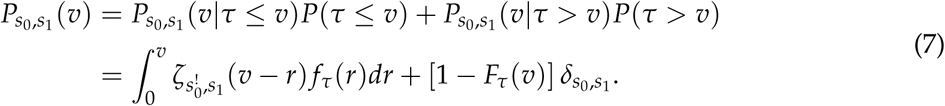

Armed with the transition probability function for our renewal model (eq. [7]), we can use the pruning algorithm to compute the likelihood of the tip state data given the tree and the model, conditional on the state at the root (Felsenstein 1981). Because we have assumed that transitions between the states are symmetric, and that the trait evolution process has been running for a long time before the root, each root state is equally probable. The full likelihood is thus the sum of the conditional likelihoods with weight one-half each.

### Simulation tests

In principle, the likelihood function derived in the previous section could be used to infer the parameters of the two-state symmetric renewal process model from phylogenetic data. To test how well this might work in practice, we implemented the likelihood calculation and used it for parameter estimation on simulated data. The limited results we report here give a rough sense of the feasibility of identifying memory in trait evolution from phylogenetic data, though they are by no means a comprehensive assessment.

For our inference model, we chose a Gamma distribution for the renewal function. The central inference question is thus whether the ‘shape’ parameter of this distribution is distinguishable from 1. If not, a Poisson model is sufficient to explain the data, and there is no evidence for memory in the macroevolution of the trait (fig. 1i). If 0 < shape < 1, memory works in the expected direction, with flips in the trait becoming more difficult the longer a state is held (fig. 1ii). If instead shape > 1, memory works in the opposite direction, with flips in the trait becoming increasingly likely (fig. 1iii).

In our testing procedure, we first simulated a large phylogeny under a simple birth-death model (500 tips, speciation rate 10× larger than extinction rate, tree scaled to a root age of 1). Then we simulated the evolution of a trait under the renewal process on that tree, using Gamma-distributed waiting times for flips of the binary trait. Our simulations and inference all assume symmetric trait evolution, with flips from *A* to *B* governed by the same distribution as flips from *B* to *A*. We then computed the likelihood of the tip state data on the tree using the likelihood function derived above, again with a Gamma distribution for the renewal function. We used Bayesian inference to estimate the shape and rate parameters of each simulation of trait evolution. We fit the model with Markov chain Monte Carlo (MCMC) using a slice sampler (Neal 2003). We assigned a prior on each parameter that was exponential with rate – ln(1/2) = 0.693, which gives equal weight to shape parameters less than or greater than 1 over the age of the tree, and which is also relatively uninformative over reasonable values of the rate parameter. To visualize how the data provide information about the shape and rate parameters, we additionally computed the likelihood on gridded parameter space. This also serves as a check that maximum likelihood parameter estimates are in general agreement with those from Bayesian inference. Our C and R code for all these procedures is included as Supplementary Material.

Our primary inference question is whether typical phylogenetic comparative data—a ‘known’ tree and trait values for extant species—bear any signal of memory in the evolution of the trait. We find that in many cases they do. Datasets simulated with a declining hazard function—so that trait flips become less likely with longer duration in a state—yielded estimates of the shape parameter that were consistently close to the true value and less than 1, though the estimates were not always precise enough to exclude 1 (fig. 6, top row). Datasets simulated with flat or increasing hazard functions yielded larger shape estimates, but these usually did not rule out a shape value of 1 with any confidence (fig. 6, middle and bottom rows). The hazard functions and rate parameter estimates are shown in figures S1–S2.

**Figure 6:**
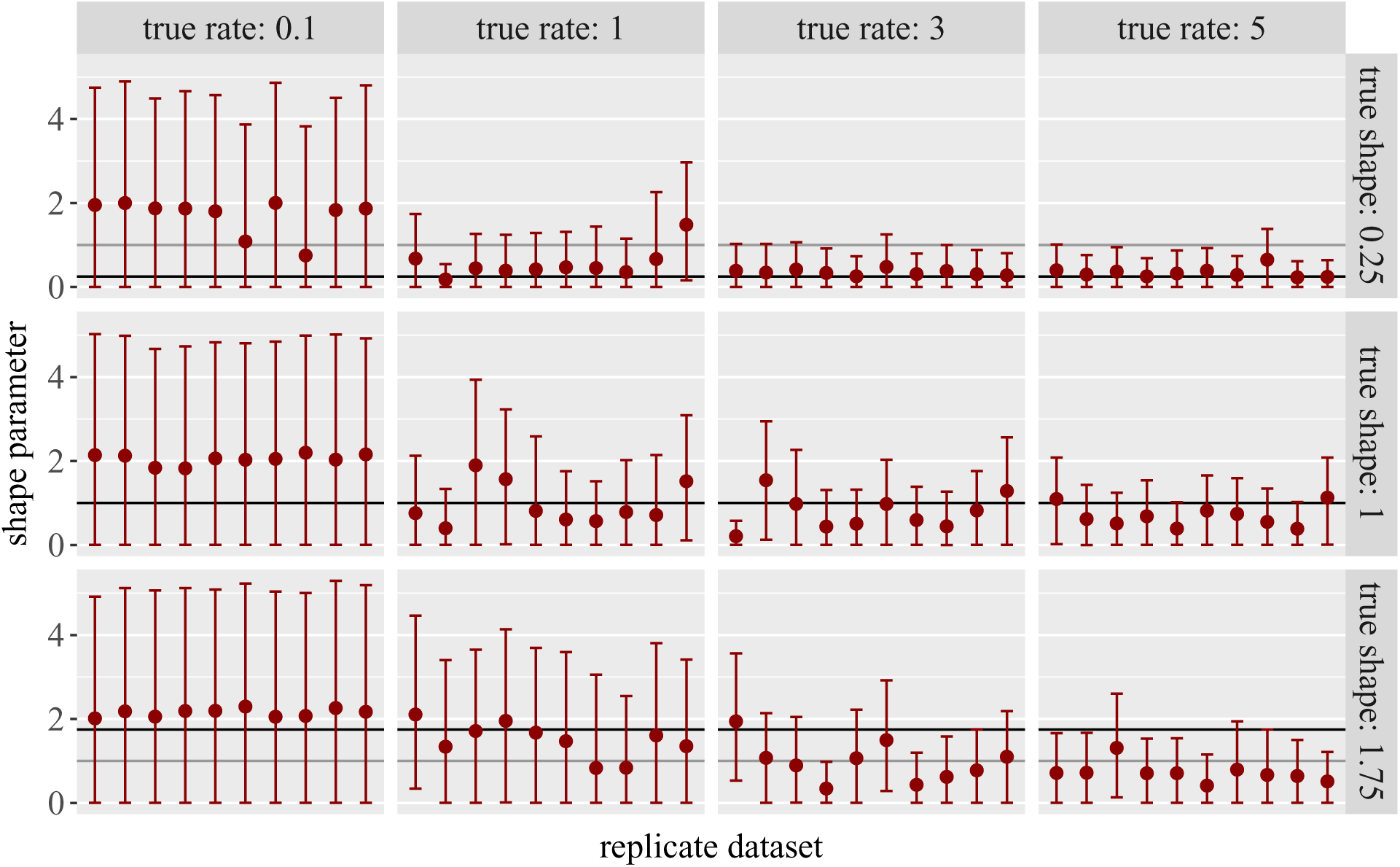
Inference results for trait evolution simulations. In each panel, results are shown for 10 datasets, each simulated on a tree with 500 tips and a root age of 1. A Gamma distribution of waiting times was used to simulate trait evolution, and its ‘shape’ and ‘rate’ parameter values are shown in the panel labels. The hazard function is either decreasing (shape of 0.25, top row), flat (shape of 1, middle row), or increasing (shape of 1.75, bottom row); these true values are marked with black horizontal lines. The full hazard functions are plotted in figure S1. The key inference question is whether the shape parameter is distinguishable from 1 (emphasized with a darker gray guide line). Inference of the shape parameter is summarized here based on the MCMC results, showing median values (points) and 90% credibility intervals (whiskers). Corresponding estimates of the rate parameter are shown in figure S2.

Estimates were less accurate and less precise when the true rate parameter was low (fig. 6, left columns). With a low rate, flips are rarer overall so less of the total branch length on the tree lies shortly after a trait flip. Because the hazard function changes most rapidly shortly after a trait flip, lower rates provide less potential to see the influence of trait duration on the instantaneous rate of change. Accuracy also appears to be worse for shape parameters larger than 1. Again, the distinguishing portion of time is shortly after a flip, but this is when the rate is low (fig. 2iii) so there are few events to inform the value of the instantaneous rate.

Visualizing the likelihood function reveals that much uncertainty comes from parameter correlations (fig. S3). There is a ridge in the likelihood surface such that the data are explained almost equally well by large shape and rate values, or by small shape and rate values. One explanation may be that the main distinguishable signal is of merely the average time between renewals, which is governed by the ratio between shape and rate parameters for the Gamma distribution choice of renewal times. For example, the three hazard functions shown in figure 1 have positively correlated parameters [shape and rate both low for (ii), both high for (iii), both intermediate for (i)] and roughly the same average value over the time interval shown. Fixing the rate parameter to the true value sidesteps the correlation and yields greatly improved estimates of the shape parameter (consider a horizontal transect in fig. S3), but this type of extra information may be difficult to obtain for real-world applications.

In summary, the Threshold, Reset, and Retain models discussed earlier provide some general guidance on the form the renewal function would take under various assumptions of the cause of memory in trait evolution. Based on that guidance, we chose one functional form for the renewal function, simulated trait evolution under it, and tested whether those simulated phylogenetic data revealed whether the true hazard function was flat, decreasing, or increasing. We found that phylogenetic comparative data do bear some signal of the shape of the hazard function, though precision and accuracy are not especially great. Thus, for future empirical studies, it may be possible to estimate the strength of memory in trait macroevolution, but further work would be needed, as discussed below.

## Discussion

Here we have considered whether trait evolution on long timescales might not be ‘memory-less,’ such that the longer a lineage has held a trait value, the harder it is for that value to change. Our goal was to describe a new macroevolutionary model of trait evolution that incorporates sufficient complexity to open up the study of this question, while retaining sufficient simplicity that it can represent evolution on many different lineages and be fit to phylogenetic data. We compared different mathematical models that incorporate memory in trait evolution, and we showed how a fairly general model can be fit to a phylogeny. We found that phylogenetic comparative data can in principle bear the signature of trait evolution memory, but that in practice there may be substantial uncertainty in the inference of this process. We end by discussing how future work might build on our approach by extending the mathematics employed, the data provided, and the questions posed.

### Extending the mathematical framework

Enhancing the mathematical models described above would open new possibilities for modeling memory in trait evolution. In some applications, the substates of the Reset or Retain model might represent known subtraits or genetic changes. If this knowledge provided more specific guidance on the difficulty of moving between substates, the transitions could be adjusted accordingly (e.g., replacing *ρ* with *ρ_i_*, or using a non-Poisson process). The allowed transitions could also be altered, to provide, for example, a mix of the Reset and Retain dynamics.

In many applications, trait evolution is expected to proceed differently in one direction than another. All of our models could be extended to accommodate this change. For the Reset and Retain models, asymmetric flips in the focal trait could be introduced by adding parameters (replacing *η_i_* with *η_Ai_* and *η_Bi_*). For the Threshold model, an asymmetric random walk could be used. For inference with a directly-chosen renewal function, the likelihood calculation could be expanded to allow an alternating renewal process.

To infer from data whether there is memory in trait macroevolution, the key inference goal is the value of the parameter that governs the presence of memory. In our simulation tests, this was the shape parameter of the Gamma distribution, but we found that its estimation was confounded with the rate parameter. To avoid this problem of parameter correlations, it might be possible to choose a different renewal distribution in which only one parameter governs the mean. Another reason to implement other functions for the renewal process is to capture hazard functions that represent different mechanisms of trait evolution. Such an extension would not require a change to the likelihood derivation, but it would require changes to the software implementation. In particular, the choice of Gamma distributed renewal times is convenient because its *n*-fold convolution, which we used in the likelihood calculation, follows a simple parametric form. A compound Poisson distribution, for example, would also possess this property. Otherwise, it may be possible to use more general classes of distributions if the *n*-fold convolution is precomputed numerically and stored for likelihood computations.

The threshold model is already in use, but its current phylogenetic applications are computationally difficult because they integrate over all the possible values of the liability at each node and tip (Felsenstein 2005; Revell 2014; Hiscott et al. 2016). Our approach is different: we work directly with the transition probabilities for the observed binary trait, not with the unobserved liabilities. Therefore, using our likelihood function with the hazard function of the threshold model, which we also computed, might provide a more efficient means of fitting the threshold model to phylogenetic data.

### Extending the data in phylogenetic comparative analyses

The simulation tests we reported are a first indication of whether one could hope to infer the presence of memory in trait macroevolution from typical phylogenetic comparative data. We find that there is indeed some signal, but that precision and accuracy may not be high. One tack for improving inference of the renewal process is to consider how other sources of information could be incorporated into an analysis.

Other studies have demonstrated that combining fossil information with phylogenetic analyses can aid inference of trait evolution (Finarelli and Flynn 2006; Slater et al. 2012; Hunt 2013; Slater 2013). We thus tested briefly whether additional information about past states might improve inference of the renewal process parameters. As an optimistic scenario, we considered the case where all species on a simulated birth-death tree are retained, whether or not they survive to the present, along with their terminal trait values. We found that on a tree with half extant tips and half extinct tips, parameter estimates were better than when the same tree was pruned to only extant tips, and that estimates were comparable to those on a different tree with the same total number of tips, all extant. (Detailed results are not shown. But more specifically, we increased the extinction rate to half the speciation rate to obtain a simulated tree with 250 extant tips and 247 extinct tips. Then we simulated the binary trait on this tree with shape = 0.25 or 1.75 and rate = 3 and used all 497 taxa for inference. We compared this to inference on the same tree pruned to the 250 extant tips, and to our main results for the same parameter values on a tree with 500 extant tips.) Thus, our brief tests indicate that fossil data do help by increasing the number of species with known state, but that the insight of extinct tips into past states does not seem to provide a particular benefit.

Besides tips representing extinct species, other kinds of historical information can anchor trait values along the branches of the tree. In the ideal case, knowing the trait values along every lineage would pinpoint the times of every trait flip and provide complete information about the renewal process. A useful next step would be to investigate whether a reasonable subset of this information on ancestral trait values could greatly improve inference of the renewal process. Even if the past trait values of a lineage cannot be precisely dated, knowing the number of trait changes over a window of time could also be helpful. Other work on renewal processes with Gamma interarrival times shows that data on the number of renewals within the time period of observation can aid parameter inference (Miller and Bhat 1997).

Even for clades with no fossil record, other kinds of information can hint at past trait values. For example, the relative degree of degeneration in underlying genes might indicate that some lineages have lost, say, functional eyes or blue flowers more recently than others (Niemiller et al. 2013; Wessinger and Rausher 2015). Such an indication of how long a lineage has held its current value of the focal trait could be incorporated by refining the binary tip state coding to the substate level in the Reset or Retain models, or perhaps by placing priors on transition times. This could potentially improve inference of the renewal process.

### Extending questions about memory in trait evolution

Our focus has been on the mathematical form and phylogenetical signal of memory in trait evolution. The models presented here may, however, also be useful in other settings.

One question in molecular evolution is whether the rate of sequence evolution depends on the state of an ecological or morphological trait (Mayrose and Otto 2011; Levy Karin et al. 2017). A renewal model could extend this question to whether the rate of sequence evolution increases after a change in the organismal trait, perhaps reflecting adaptation that is most rapid initially. For example, one could use standard Poisson models for the organismal-level trait and for sequence evolution, but additionally with the overall rate of base pair change following a renewal process, based on the time since the last organismal trait flip. Such an application is likely to derive much more power from the many sites in a sequence: each site evolves under the same model, and all have the same rate at a given time.

The memory model of trait evolution could also be coupled with models of lineage diversification. For example, increasing inability to adapt to a shift in selective regime could result in duration-dependent extinction. This resembles the model of Alexander et al. (2016), but the critical factor is time since the last trait change rather than time since the lineage’s origination. An implementation would involve replacing transition probabilities with differential equations for clade and extinction probabilities (as in Maddison et al. 2007).

An initial motivation in developing the renewal model of trait evolution was that it might alleviate problems of phylogenetic pseudoreplication in studying trait evolution. For testing correlations between two discrete-valued traits, or between one trait and lineage diversification rates, existing methods draw ‘signal’ from all parts of the tree that exhibit the correlation, instead of from the number of independent times that association has arisen (Maddison and FitzJohn 2015; Rabosky and Goldberg 2015). Perhaps a trait evolution model in which the time since the last change plays an important role would be less susceptible to this problem.

Finally, we will be curious to see if this approach to modeling trait evolution has utility in other areas of ecology and evolution. For example, consider a theoretical investigation of when competitors can coexist on resources that change with time. A renewal process could capture the idea that the longer one participant has specialized on a single resource, the harder it is to switch to another. The coexistence dynamics of such a model might differ from formulations with other inhibitions to resource switching.

## Conclusion

Our premise has been that the longer a lineage holds a trait value, the harder may become evolution away from that value. This is, however, only a hypothesis. Evolution does indeed take time, but whether the ‘memory’ dynamic of trait evolution emerges at a macroevolutionary scale depends on how elapsed time relates to extent of fit with the environment, and the degree to which increased fit to one regime inhibits evolution in a new direction. We hope that the present work will enable broad comparative tests that complement system-specific investigations of these questions.

## Acknowledgements

We are grateful to Maria Servedio for organizing the symposium and imposing deadlines that enforced progress on this project. We thank members of the UMN EEB ‘Theory under construction’ group for their comments in the early stages. Tanjona Ramiadantsoa implemented early simulations of the renewal process. Will Freyman suggested testing inference with fossil tips and applying the renewal model to molecular sequence evolution. This work was supported by National Science Foundation grants DEB-1655478 to EEG and DMS-1349724 to JF.

---

**Figure S1:**
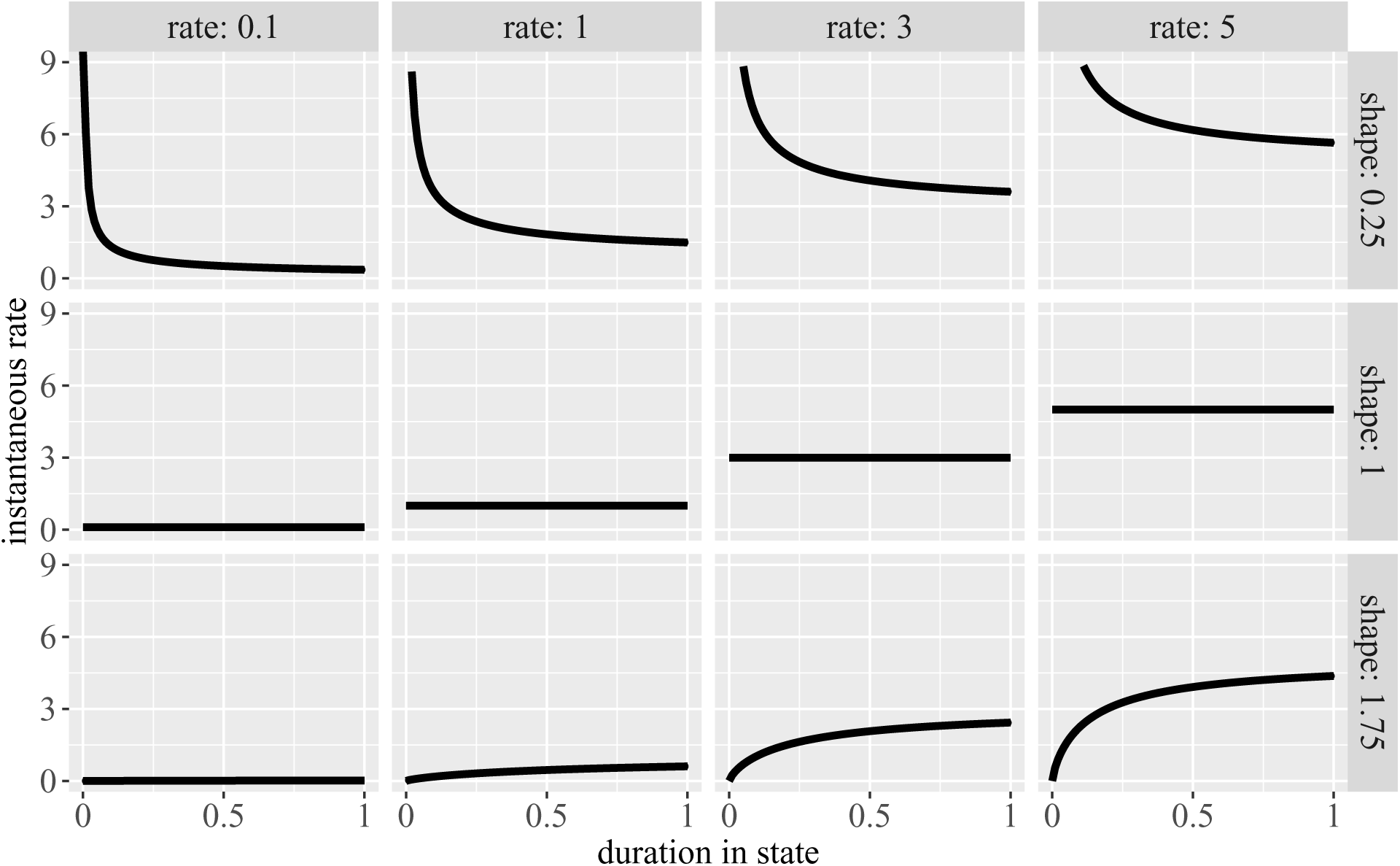
Hazard functions used for simulation tests reported in figure 6 and figure S2.

**Figure S2:**
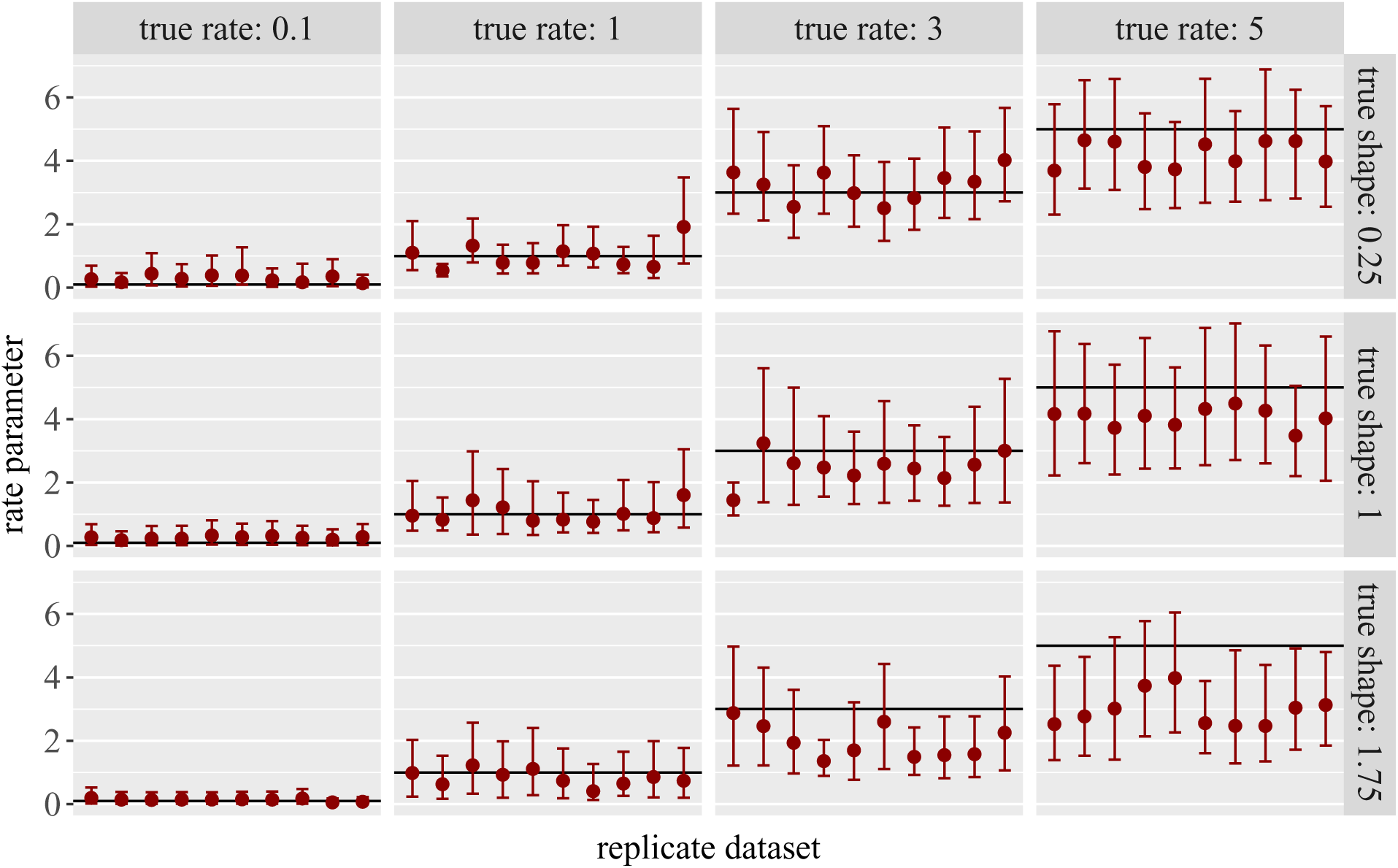
More inference results for trait evolution simulations. For the same simulated datasets, estimates of the shape parameter are shown in figure 6 and estimates of the rate parameter are shown here.

**Figure S3:**
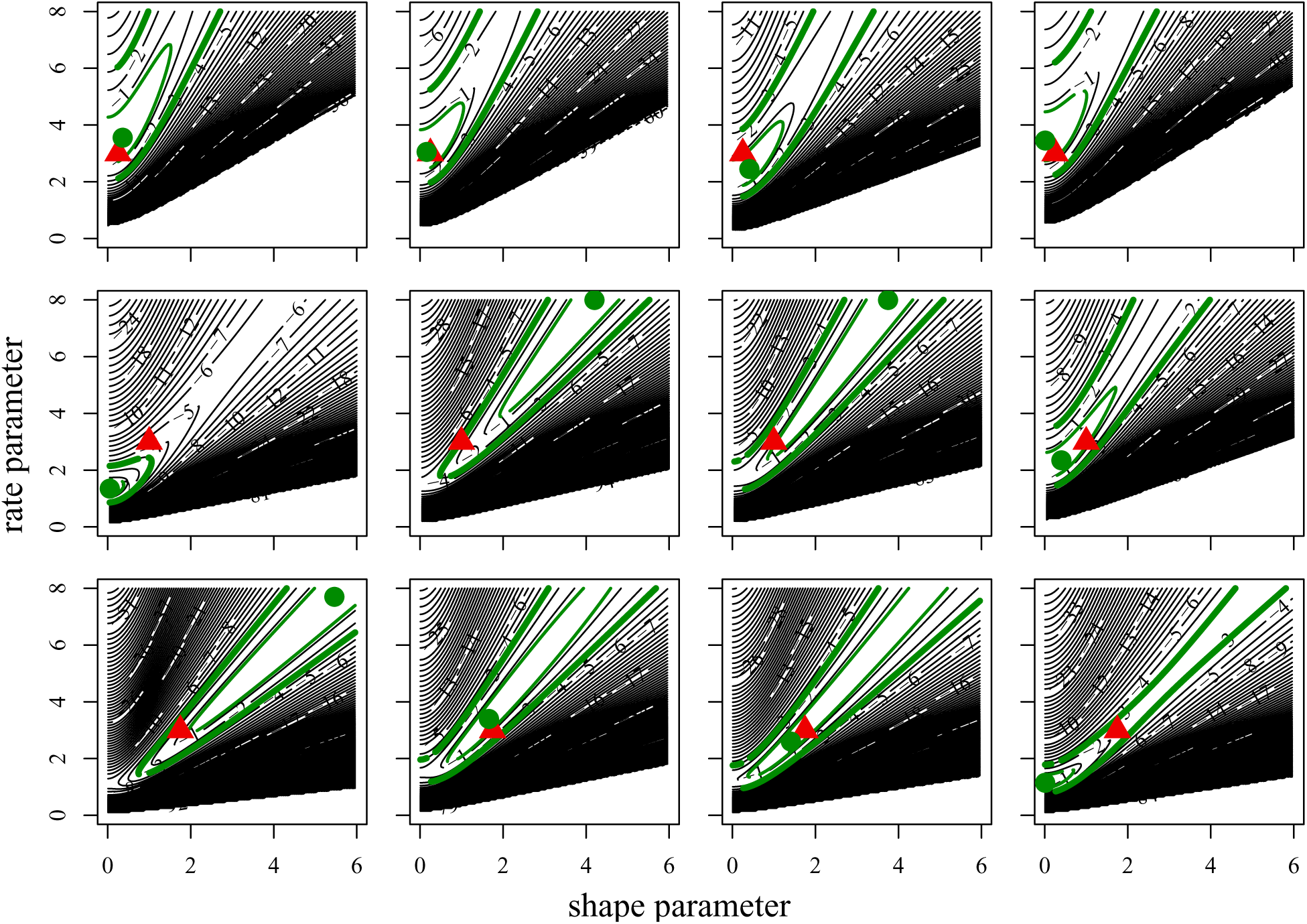
Log-likelihood surfaces for ten simulated datasets on trees with 500 tips. Datasets are the first four for each shape value, with a rate value of 3, in figure 6 and figure S2. True parameter values are marked with red triangles. Maximum likelihood estimates are marked with green circles. Black contour line spacing is 1 log-likelihood unit, and the log-likelihood values are normalized so that the maximum is 0. Green contours additionally mark the 50% and 95% likelihood ratio confidence intervals, computed with the chi-squared approximation.

## Literature Cited

Alexander, H. K., A. Lambert, and T. Stadler. 2016. Quantifying age-dependent extinction from species phylogenies. Systematic Biology 65:35–50.

Beaulieu, J. M., and B. C. O’Meara. 2016. Detecting hidden diversification shifts in models of trait-dependent speciation and extinction. Systematic Biology 65:583–601.

Beaulieu, J. M., B. C. O’Meara, and M. J. Donoghue. 2013. Identifying hidden rate changes in the evolution of a binary morphological character: the evolution of plant habit in campanulid angiosperms. Systematic Biology 62:725–737.

Bokma, F. 2008. Detection of “punctuated equilibrium” by Bayesian estimation of speciation and extinction rates, ancestral character states, and rates of anagenetic and cladogenetic evolution on a molecular phylogeny. Evolution 62:2718–2726.

Bull, J. J., and E. L. Charnov. 1985. On irreversible evolution. Evolution 39:1149–1155.

Charlesworth, D. 2015. Plant contributions to our understanding of sex chromosome evolution. New Phytologist 208:52–65.

Felsenstein, J. 1981. Evolutionary trees from DNA sequences: a maximum likelihood approach. Journal of Molecular Evolution 17:368–376.

Felsenstein, J.. 2005. Using the quantitative genetic threshold model for inferences between and within species. Philosophical Transactions of the Royal Society B: Biological Sciences 360:1427–1434.

Finarelli, J. A., and J. J. Flynn. 2006. Ancestral state reconstruction of body size in the Caniformia (Carnivora, Mammalia): the effects of incorporating data from the fossil record. Systematic Biology 55:301–313.

Freyman, W., and S. Höhna. in review. The tempo of evolutionary decline in self-compatible plant lineages. bioRxiv

Goldberg, E. E., and B. Igić. 2008. On phylogenetic tests of irreversible evolution. Evolution 62:2727–2741.

Goldberg, E. E., and B. Igić. 2012. Tempo and mode in plant breeding system evolution. Evolution 66:3701–3709.

Goldberg, E. E., S. P. Otto, J. C. Vamosi, I. Mayrose, N. Sabath, R. Ming, and T.-L. Ashman. 2017. Macroevolutionary synthesis of flowering plant sexual systems. Evolution 71:898–912.

Goldman, N., and Z. Yang. 1994. A codon-based model of nucleotide substitution for protein-coding DNA sequences. Molecular Biology and Evolution 11:725–736.

Hagen, O., K. Hartmann, M. Steel, and T. Stadler. 2015. Age-dependent speciation can explain the shape of empirical phylogenies. Systematic Biology 64:432–440.

Hiscott, G., C. Fox, M. Parry, and D. Bryant. 2016. Efficient recycled algorithms for quantitative trait models on phylogenies. Genome Biology and Evolution 8:1338–1350.

Hunt, G. 2013. Testing the link between phenotypic evolution and speciation: an integrated palaeontological and phylogenetic analysis. Methods in Ecology and Evolution 4:714–723.

Lalley, S. 2016. Random walk lecture notes.

Levy Karin, E., S. Wicke, T. Pupko, and I. Mayrose. 2017. An integrated model of phenotypic trait changes and site-specific sequence evolution. Systematic Biology 66:917–933.

Lewis, P. O. 2001. A likelihood approach to estimating phylogeny from discrete morphological character data. Systematic Biology 50:913–925.

Maddison, W. P., and R. G. FitzJohn. 2015. The unsolved challenge to phylogenetic correlation tests for categorical characters. Systematic Biology 64:127–136.

Maddison, W. P., P. E. Midford, and S. P. Otto. 2007. Estimating a binary character’s effect on speciation and extinction. Systematic Biology 56:701–710.

Magnuson-Ford, K., and S. P. Otto. 2012. Linking the investigations of character evolution and species diversification. The American Naturalist 180:225–245.

Marshall, C. R., E. C. Raff, and R. A. Raff. 1994. Dollo’s law and the death and resurrection of genes. Proceedings of the National Academy of Sciences 91:12283–12287.

Mayrose, I., and S. P. Otto. 2011. A likelihood method for detecting trait-dependent shifts in the rate of molecular evolution. Molecular Biology and Evolution 28:759–770.

McGaugh, S. E., J. B. Gross, B. Aken, M. Blin, R. Borowsky, D. Chalopin, H. Hinaux, W. R. Jeffery, A. Keene, L. Ma, P. Minx, D. Murphy, K. E. O’Quin, S. Rétaux, N. Rohner, S. M. J. Searle, B. A. Stahl, C. Tabin, J.-N. Volff, M. Yoshizawa, and W. C. Warren. 2014. The cavefish genome reveals candidate genes for eye loss. Nature Communications 5:5307.

Miller, G., and U. N. Bhat. 1997. Estimation for renewal processes with unobservable gamma or Erlang interarrival times. Journal of Statistical Planning and Inference 61:355–372.

Neal, R. M. 2003. Slice sampling. Annals of Statistics 31:705–741.

Niemiller, M. L., B. M. Fitzpatrick, P. Shah, L. Schmitz, and T. J. Near. 2013. Evidence for repeated loss of selective constraint in rhodopsin of amblyopsid cavefishes (Teleostei: Amblyopsidae). Evolution 67:732–748.

Nosil, P., and A. Ø. Mooers. 2005. Testing hypotheses about ecological specialization using phylogenetic trees. Evolution 59:2256–2263.

O’Meara, B. C., C. Ané, M. J. Sanderson, and P. C. Wainwright. 2006. Testing for different rates of continuous trait evolution using likelihood. Evolution 60:922–933.

Pagel, M. 1994. Detecting correlated evolution on phylogenies: a general method for the comparative analysis of discrete characters. Proceedings of the Royal Society of London, Series B 255:37–45.

Rabosky, D. L., and E. E. Goldberg. 2015. Model inadequacy and mistaken inferences of trait-dependent speciation. Systematic Biology 64:340–355.

Ree, R. H., B. R. Moore, C. O. Webb, and M. J. Donoghue. 2005. A likelihood framework for inferring the evolution of geographic range on phylogenetic trees. Evolution 59:2299–2311.

Revell, L. J. 2014. Ancestral character estimation under the threshold model from quantitative genetics. Evolution 68:743–759.

Ross, S. M. 2010. Introduction to Probability Models. 10th ed. Academic Press.

Slater, G. J. 2013. Phylogenetic evidence for a shift in the mode of mammalian body size evolution at the Cretaceous-Palaeogene boundary. Methods in Ecology and Evolution 4:734–744.

Slater, G. J., L. J. Harmon, and M. E. Alfaro. 2012. Integrating fossils with molecular phylogenies improves inference of trait evolution. Evolution 66:3931–3944.

Stadler, T. 2013. Recovering speciation and extinction dynamics based on phylogenies. Journal of Evolutionary Biology 26:1203–1219.

Tarasov, S. in review. Integration of anatomy ontologies and evo-devo using structured Markov models suggests a new framework for modeling discrete phenotypic traits. bioRxiv.

Wessinger, C. A., and M. D. Rausher. 2015. Ecological transition predictably associated with gene degeneration. Molecular Biology and Evolution 32:347–354.

Whittall, J. B., and S. A. Hodges. 2007. Pollinator shifts drive increasingly long nectar spurs in columbine flowers. Nature 447:706–709.

Wright, S. 1934. The results of crosses between inbred strains of guinea pigs, differing in number of digits. Genetics 19:537–551.

Zenil-Ferguson, R., J. M. Ponciano, and J. G. Burleigh. 2017. Testing the association of phenotypes with polyploidy: An example using herbaceous and woody eudicots. Evolution 71:1138–1148.

